# Network Enrichment Significance Testing in Brain-Phenotype Association Studies

**DOI:** 10.1101/2023.11.10.566593

**Authors:** Sarah M. Weinstein, Simon N. Vandekar, Aaron F. Alexander-Bloch, Armin Raznahan, Mingyao Li, Raquel E. Gur, Ruben C. Gur, David R. Roalf, Min Tae M. Park, Mallar Chakravarty, Erica B. Baller, Kristin A. Linn, Theodore D. Satterthwaite, Russell T. Shinohara

**Author notes:** Corresponding author: S.M. Weinstein.

## Abstract

Functional networks often guide our interpretation of spatial maps of brain-phenotype associations. However, methods for assessing enrichment of associations within networks of interest have varied in terms of both scientific rigor and underlying assumptions. While some approaches have relied on subjective interpretations, others have made unrealistic assumptions about the spatial structure of imaging data, leading to inflated false positive rates. We seek to address this gap in existing methodology by borrowing insight from a method widely used in genomics research for testing enrichment of associations between a set of genes and a phenotype of interest. We propose Network Enrichment Significance Testing (NEST), a flexible framework for testing the specificity of brain-phenotype associations to functional networks or other sub-regions of the brain. We apply NEST to study phenotype associations with structural and functional brain imaging data from a large-scale neurodevelopmental cohort study.

## 1 Introduction

Quantifying and spatially mapping brain-phenotype associations is a central component of many neuroimaging studies. There is often particular interest in interpreting these maps through the lens of canonical functional networks, which delineate areas of the brain known to participate in different behaviors and cognitive functions (Yeo et al. 2011; Schaefer et al. 2018). Evaluating whether strong brain-phenotype associations are especially strong—or *enriched* —within a network of interest can add to our understanding of the neural mechanisms underlying transdiagnostic psychopathology (Segal et al. 2023). However, the methodologies underlying previous claims about network enrichment have varied and even relied on subjective interpretations—for instance, visualizing a map of brain-phenotype associations next to a network parcellation map and noting any networks with which strong associations appear to coincide. While such claims are often compelling, the reliance on subjective interpretations, rather than statistical methods with well-defined null hypotheses, may compromise replicability.

Some researchers have indeed shifted towards conducting more reproducible tests examining spatial overlap between local brain-phenotype associations and functional networks. Alexander-Bloch et al. (2018)’s *spin test*, which evaluates spatial alignment between two maps, has become especially popular. Essentially, the spin test offers a procedure by which to test spatial alignment between a map of brain-phenotype associations to a network map (e.g., Figure 1(A) to (B)). Briefly, the analyst first computes a correlation (or other similarity metric) between the two maps, then projects one of the two maps onto a sphere, randomly rotates (or “spins”) it, and computes the correlation again. In a procedure analogous to permutation testing, the analyst repeats these random rotations, and the correlations computed from the “spun” data form a null distribution, which is compared to the original correlation to estimate a *p*-value. The spin test has been used to test claims about network alignment with maps of brain-phenotype associations and inter-modal coupling (Mandal et al. 2020; Baum et al. 2020; Nadig et al. 2021; Petersen et al. 2022; Baller et al. 2022; Hu et al. 2022; Markello et al. 2022). Several adaptations and extensions of the method have also been described (Váša et al. 2018; Vázquez-Rodríguez et al. 2019; Cornblath et al. 2020).

**Figure 1.**
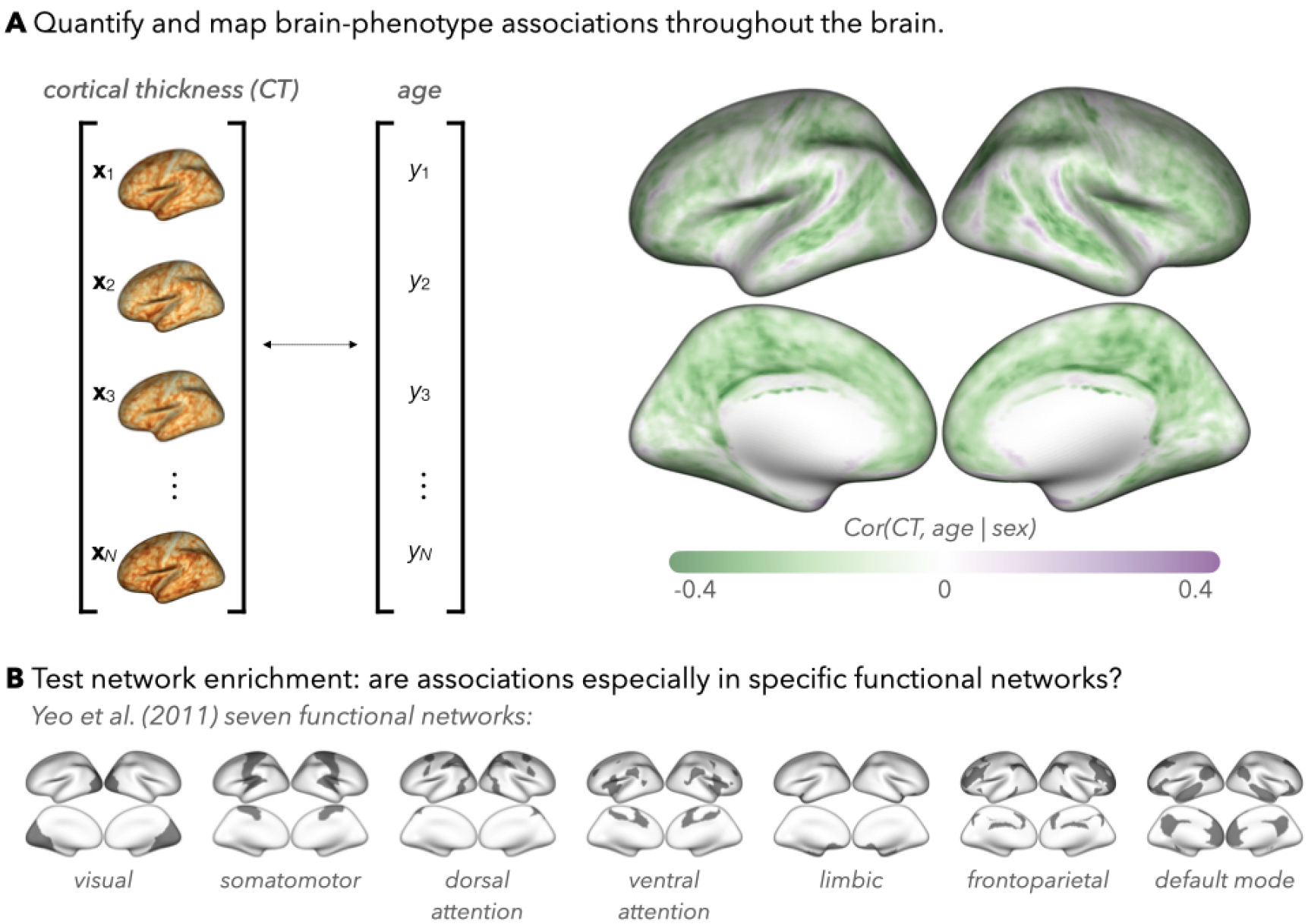
Example of how brain-phenotype associations are (**A**) quantified and spatially mapped and **(B)** compared to maps of functional networks (e.g., those delineated by Yeo et al. (2011)). As we discuss in Section 1, existing methods for evaluating network specificity (or “enrichment”) of brain-phenotype associations have been subjective, which may preclude reproducibility, or relied on strong assumptions, which may result in type I error inflation. In Section 2, we propose a new approach, called NEST, to address these limitations.

Clearly, the spin test is an important step forward, setting a more rigorous methodological standard for brain map comparisons (Burt et al. 2020; Markello and Misic 2021; Weinstein et al. 2021; Weinstein et al. 2022). However, it is unclear whether the method’s underlying null hypothesis is an appropriate translation of many researchers’ scientific hypotheses concerning network enrichment—that is, whether associations are especially strong within a network of interest. In particular, the spin test evaluates spatial alignment between two maps, which does not necessarily correspond to stronger associations being found within the network. It is plausible, for example, that the spin test might yield a statistically significant result if an association map contained greater spatial smoothness—but not necessarily stronger associations—within networks compared to between networks. Finally, several limitations of the spin test have been noted, including that it can only be applied to cortical surface data and that it requires assuming covariance stationarity for statistical inference to be generalizable across datasets (Burt et al. 2020; Markello and Misic 2021; Weinstein et al. 2021). Further exploration of these limitations and of alternative methods is warranted.

Another approach was described by Park et al. (2018) in a study of enrichment of associations between brain structure and neuropsychiatric diagnoses. These authors borrowed insight from Gene Set Enrichment Analysis (GSEA) (Subramanian et al. 2005), which has been widely used in genomics research to test enrichment of sets of genes for particular phenotypes—for example, to examine the collective role of a group of genes (rather than individual genes) on disease risk. They specifically adopted a variation of GSEA, known as FastGSEA (Korotkevich et al. 2016). Both versions of the method involve computing an “enrichment score,” which quantifies the relative strength of gene-phenotype associations for genes found within versus outside a pre-specified set of genes, using permutations to derive a null distribution. The main difference between these two approaches lies in the unit of permutation: while GSEA permutes samples before re-computing the enrichment score to form a null distribution, FastGSEA permutes gene labels. The translation of FastGSEA to neuroimaging involves permuting network labels in a manner that destroys the underlying spatial structure of the data, violating the exchangeability assumption essential for type I error control in permutation tests. In this paper, we propose Network Enrichment Significance Testing (NEST, adapting GSEA by Subramanian et al. (2005)). The proposed method operationalizes the notion of enrichment through a test statistic which quantifies the degree to which associations found inside the network are “more extreme” than those found outside the network. As we will illustrate, NEST controls type I error rates by incorporating a permutation-based null distribution that does not make assumptions about the spatial structure of the data. The remainder of this article is organized as follows. In Section 2, we define the underlying null hypothesis and implementation of our proposed method. In Section 3, we apply and compare our method with previous approaches in studies of network enrichment of brain-phenotype associations using structural and functional MRI data from the Philadelphia Neurodevelopmental Cohort. Finally, in Section 4, we discuss implications, limitations, and future directions of our method.

## 2 Methods

### 2.1 Null hypothesis

Let *T* (*v*) denote a statistic of brain-phenotype associations (e.g., correlation or a metric derived from a regression model), computed for a given image location *v* (*v* = 1, …, *V*), using data across *N* participants (e.g., as in Figure 1(A)). Let *𝒩* ⊆ {1, …, *V*} denote a subset of all vertices indexing a network of interest (e.g., from Figure 1(B)). Let 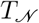 denote a random variable whose realizations are *T* (*v*) from randomly selected locations inside the network (*T* (*v*) for *v* ∈ *𝒩*), and let 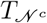 denote a random variable whose realizations are found at locations outside the network (*T* (*v*) for *v ∉ 𝒩*). We conceptualize network enrichment as any setting in which realizations of *T*_*𝒩*_ are more extreme (i.e., either more strongly positive or more strongly negative) than realizations of 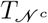, and we propose evaluating enrichment by comparing the stochastic ordering of these random variables.

For two random variables *Y* and *Z*, if

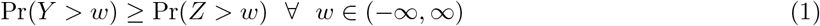

then *Y* is stochastically greater than *Z* (or *Y ⪰*_*st*_ *Z*) and “strictly” stochastically greater (or *Y ≻*_*st*_ *Z*) if there exists a *w* such that Pr(*Y > w*) *>* Pr(*Z > w*) (Belzunce et al. 2015). We consider a variation of this concept for the current setting, where we are interested in characterizing stochastic ordering in terms of the “extremeness” of realizations of the *T*_*𝒩*_ and 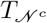 . Here, we characterize the random variable *Y* to be “stochastically more extreme” than *Z* if either of the following is true:

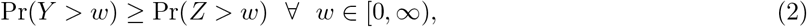

Or

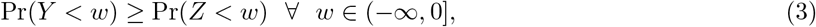

and additionally, if either (2) or (3) holds strictly for some *w*. We denote this relationship with the following notation:

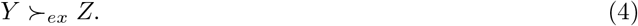

Finally, our proposed null hypothesis for testing enrichment in a given network *𝒩* is

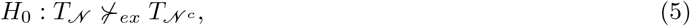

so that when *H*_0_ is true, *T*_*𝒩*_ is “stochastically no more extreme” than 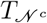 . Under the alternative hypothesis, 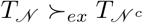.

As we will discuss in the next section, to test the proposed *H*_0_, we use a test statistic called an enrichment score (ES), which quantifies the degree to which realizations of *T*_*𝒩*_ are more extreme than realizations of 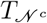. Under the alternative hypothesis, where realizations of *T*_*𝒩*_ and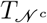 would generally be quite different (in terms of both sign and magnitude), the larger the ES would be. Under the null hypothesis, realizations of *T*_*𝒩*_ and 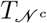 would be more similar, and the ES would be smaller.

### 2.2 Network Enrichment Significance Test (NEST)

For a given brain-phenotype association (e.g., cortical thickness, age) and pre-specified set of network labels *𝒩 ⊆* {1, …, *V*}, we propose the following adaptation of Subramanian et al. (2005)’s GSEA for 𝒩etwork Enrichment Significance Testing (NEST).

1. Quantify brain-phenotype associations *T* (*v*) at each location (*v* = 1, …, *V*). The choice of metric is up to the researcher (e.g., partial correlation, as in Figure 1(B), regression coefficient estimates, Wald statistic, or something else), but it must have a sign.
2. Sort the *T* (*v*) in descending order, so that the *v*’s with the strongest positive brain-phenotype associations appear at the top of the list, and vertices with strongest negative associations appear at the bottom of the list (Figure 2(A)). In the ranked list, we denote the *j*th entry as *T* (*v*_(*j*)_) = (equal to the *j*th largest *T* (*v*) across all *v*’s), where the subscript *j* = 1, …, *V* is the rank and *v* as well as *v*_(*j*)_ still denote the original vertex label, which maps back to a specific location in the brain map.

**Figure 2.**
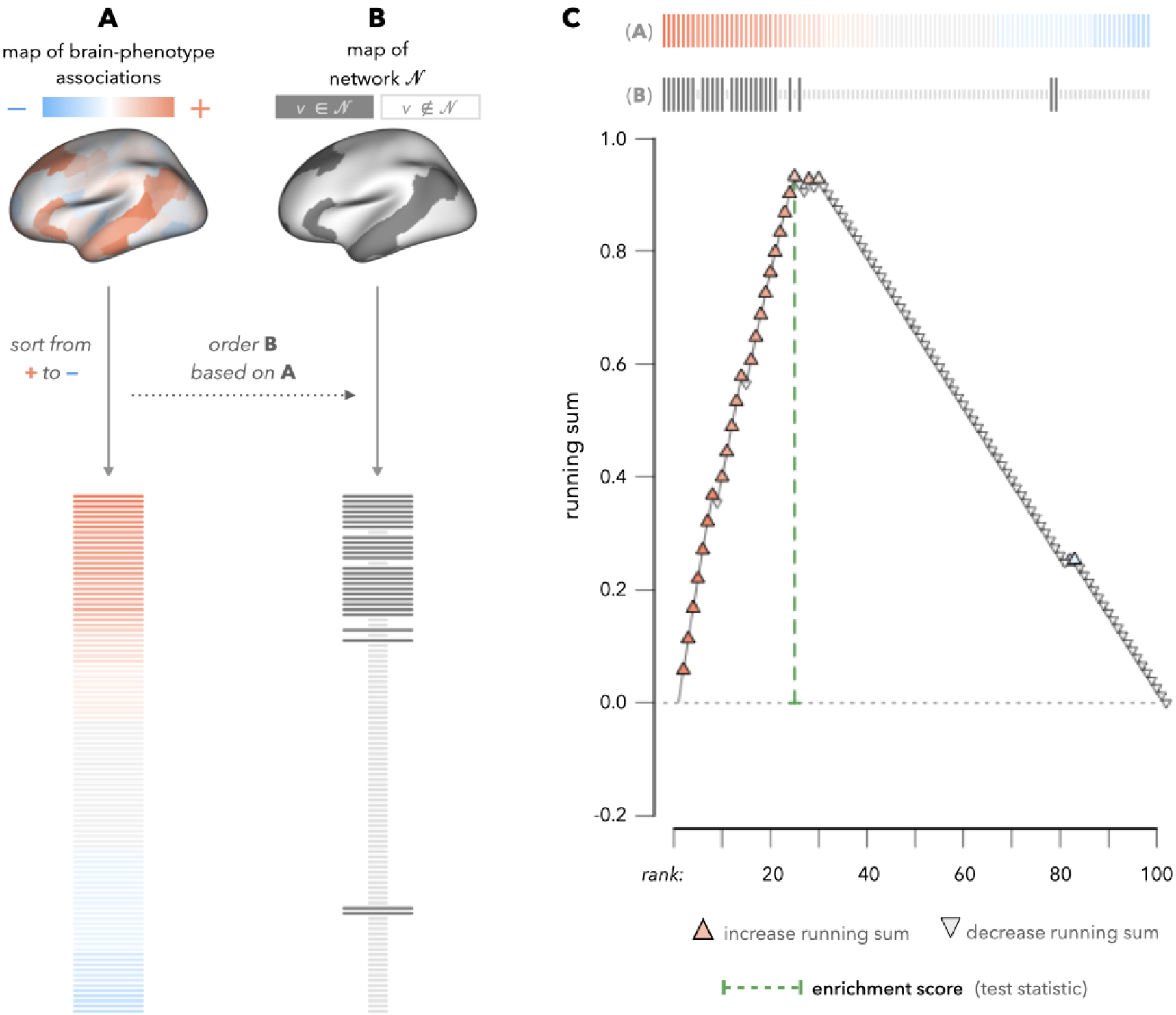
Illustration of our proposed method, NEST, which adapts Subramanian et al. (2005)’s GSEA in order to test network enrichment of brain-phenotype associations. For the purpose of this illustration, we use a simulated example of a left hemisphere fsaverage5 map using the Schaefer et al. (2018) parcellation involving 100 locations per hemisphere. (In contrast, the real data analysis described in this paper includes over 10,000 locations (vertices) per hemisphere; see Section 2.3). In extending GSEA to the neuroimaging setting, our goal is to statistically test whether brain-phenotype associations are especially strong within a network of interest, *𝒩* (binary map shown in **B** uses Yeo et al. (2011)’s default mode network as an example, resampled to the Schaefer et al. (2018) atlas with a lower dimension). In **A**, we begin with a quantitative map of brain-phenotype associations, where positive associations are represented in red and negative associations are represented in blue. Next, we sort the values of the association metric in **A** from positive to negative. In **B**, we match the order of the sorted list of association metrics in **A** to the binary brain map corresponding to *𝒩* (dark lines correspond to locations *v* ∈ *𝒩* ; faint lines correspond to locations *v ∉ 𝒩*). In **C**, we quantify the extent to which values of *T* (*v*) with larger magnitudes (i.e., either darker reds or darker blues) tend to appear within versus outside *𝒩* . A running sum statistic is initialized at 0 and increases by an increment proportional to *T* (*v*) for *v* ∈ *𝒩* . We indicate this with upward-pointing triangles, shaded based on the corresponding location in **A**. For *v* ∉ *𝒩*, we decrease the running sum statistic by a uniform increment (indicated with upside-down triangles). The enrichment score (ES, green dotted line) is the largest magnitude attained of the running sum and is used as our test statistic for evaluating *H*_0_ (Equation (5)). For inference, we repeatedly permute the phenotype of interest across participants, re-calculate the association map based on that permuted phenotype, and repeat **A**-**C**. The enrichment scores obtained from permuted data form a null distribution for testing *H*_0_.
3. In the sorted list of brain-phenotype associations, we identify the *T* (*v*_(*j*)_)’s where *v*_(*j*)_ ∈ *𝒩* (see horizontal lines in Figure 2(B)).
4. Initialize a running sum metric for network *𝒩* at 0 (*RS*_(0)_ = 0). Begin walking down the list for *j* = 1, …, *V* (i.e. starting with *T* (*v*_(1)_), the strongest positive association, and ending with *T* (*v*_(*V*)_), the strongest negative association. Increase or decrease the running sum as follows (and as illustrated in Figure 2(C)):

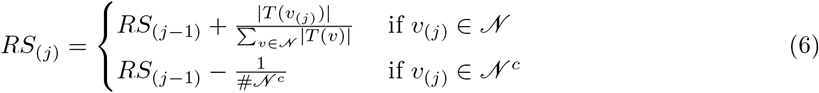

where the cardinality of the set *𝒩* ^*c*^ is denoted by #*𝒩* ^*c*^ (i.e., the number of locations outside *𝒩*).
5. Compute the enrichment score (*ES*) for network *𝒩* as the maximum deviation from zero that *RS* takes on.

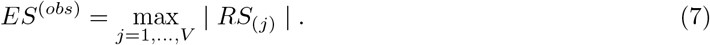

Importantly, by increasing the RS by an increment proportional to | *T* (*v*_(*j*)_) | in the previous step, the ES quantifies the extent to which associations are both stronger and of the same sign within the network compared to outside of it. (An unweighted version of the first case in Equation (6) would correspond to a Kolmogrov-Smirnov statistic.) If we were to use an unweighted statistic, we would be testing for differences in the distribution, rather than testing for stochastic extremeness as described in Section 2.1.
6. For *k* = 1, …, *K* permutations (e.g., *K* = 999), permute the phenotype across individuals and quantify brain-phenotype associations *T* (*v*) at each location in the permuted sample. Repeat steps 2-5, denoting the ES from the *k*th permutation by *ES*^(*k*)^.
7. Estimate a *p*-value for evaluating enrichment of brain-phenotype associations in network *𝒩*, by comparing *ES*^(*obs*)^ to *ES*^(*k*)^, *k* = 1, …, *K* (distribution of *ES*^(*obs*)^ under *H*_0_):

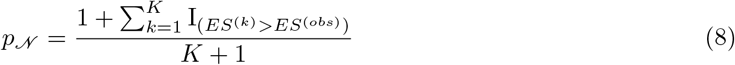

where the smallest possible *p*-value is 1*/*(*K* + 1) (Phipson and Smyth 2010).

While the steps above return a *p*-value for a pre-specified set of network labels *𝒩*, some steps do not need to be repeated if enrichment is being tested in multiple networks. Specifically, steps 1-2 remain the same across different networks (as does step 6 each time steps 1-2 are reapplied in permuted data). Additionally, we note that the specific approach to permutation (step 6) may depend presence of confounders or other nuisance variables. This topic has been explored in previous literature, and we refer readers to work by Winkler et al. (2014).

An R package for implementing NEST is currently under development and will be made available at the time of publication.

### 2.3 Data-driven simulations and data analysis in the PNC

We evaluate the performance of NEST using data from the Philadelphia Neurodevelopmental Cohort (PNC), a large-scale study of children, adolescents, and young adults between 8 and 21 years of age (Satterthwaite et al. 2016). A subset of PNC participants underwent multi-modal neuroimaging at the Hospital of the University of Pennsylvania in the same Siemens TIM Trio 3T scanner under the same protocols, which are detailed in Satterthwaite et al. (2014) and summarized in Appendix A. Utilizing the same processed data from our earlier work (Weinstein et al. 2021), data were processed using FreeSurfer version 5.3 and the fsaverage5 atlas to reconstruct images on the cortical surface, consisting of 10, 242 vertices per hemisphere (Fischl 2012). After removing the medial wall (i.e., subtracting 888 and 881 vertices from the left and right hemispheres, respectively), *V* = 18, 715 vertices remain across both hemispheres for each person. While the medial wall is excluded in our models of brain-phenotype associations, we add these vertices back for the purpose of visualizations using the `fsbrain` package in R (Schäfer and Ecker 2020).

In this study, we focus on network enrichment of age and sex effects on brain structure (cortical thickness in mm) for *𝒩* = 911 participants and functional activation during the *n*-back task (percent change between the 0- and 2-back sequences) for *N* = 1, 018 participants, after applying similar exclusion criteria as in our previous work (Weinstein et al. 2021). In simulation studies and data analyses (Sections 2.3.2 and 2.3.3 below), we apply NEST to evaluate enrichment of brain-phenotype associations in the seven functional networks described by Yeo et al. (2011). We anticipate that age and sex effects may be especially enriched within the dorsal attention, ventral attention, frontoparietal, and default networks, as increased connectivity within and between these networks play an important role in the maturation of cognitive abilities from childhood to adolescence (Sydnor et al. 2021; Keller et al. 2022). Age-related differences in brain activity during the *n*-back task are often believed to localize within these networks that are involved in higher-order functions (Satterthwaite et al. 2013). Applying NEST in analyses of brain-phenotype associations within these networks in a neurodevelopmental cohort may indeed corroborate past findings, while invoking more rigorous methodology.

#### 2.3.1 Quantifying brain-phenotype associations

In this section, we describe our approach to quantifying local linear and nonlinear brain-phenotype associations in the P𝒩C data. To flexibly model brain image measurements as a function of non-imaging phenotypes, we fit a generalized additive model (GAMs) at each vertex. We then use a multivariate Wald statistic to jointly capture linear and nonlinear brain-phenotype associations from each GAM.

Let **x**(*v*) = (*x*_1_(*v*) *x*_2_(*v*) … *x*_*N*_ (*v*)) denote measurements of the brain (across participants) at a given location *v*. We consider the following model at each vertex *v* and for each type of brain measurement (cortical thickness or *n*-back activation):

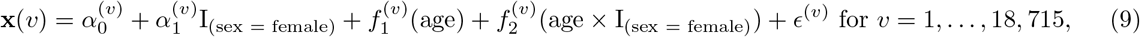

where the superscripts (*v*) indicate location (vertex)-specific components of the model, and I_()_ denotes an indicator function. In the model above, 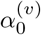 and 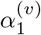 are linear parameters for mean and sex (female) effects, respectively, for each *v*. The smoothing functions, 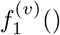 and 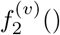, are fixed degrees of freedom regression natural cubic splines with *K*_1_ and *K*_2_ knots, respectively. We estimate these functions using the mgcv package in R (Wood and Wood 2015).

The use of these smooth functions means that we estimate multiple different parameters for each effect of interest. To spatially map brain-phenotype associations, we would still like to collapse information across these parameters into a single vertex-level statistic, *T* (*v*). To combine information across all parameters associated with each effect of interest in Equation (9), we use a multivariate Wald statistic. Since the Wald statistic is expressed as a quadratic form and will always be positive, we assign a the sign of each Wald statistic based on coefficient estimates from multiple linear regression models, which would reflect the overall trend for nonlinear associations. Details on this statistic are outlined in Appendix B.

#### 2.3.2 Simulation studies

We use a resampling procedure to conduct simulation studies using real data from the P𝒩C. Across 1,000 simulations, we consider scenarios involving sub-samples of different sizes (*N* _*sub*_ = 50, 100, 200, or 300) and evaluate whether each brain-phenotype association of interest is enriched within each of the seven functional networks delineated by Yeo et al. (2011). We conduct these simulations in both a simulated null setting (for estimating type I error) and observed data setting (for estimating power).

##### Type I error simulations

We first evaluate the type I error rates of NEST and two existing approaches: the spin test (Alexander-Bloch et al. 2018) and FastGSEA (Korotkevich et al. 2016; Park et al. 2018). For a given brain-phenotype association (e.g., cortical thickness and age), we randomly permute the phenotype across participants in the full sample, then randomly select a subset of *N*_*sub*_ (50, 100, 200, or 300). By permuting the data before subsetting, we force *H*_0_ to be true, since brain-phenotype associations have been destroyed at all locations, and thus there should be no enrichment. Even in this null setting, we nevertheless anticipate that maps of brain-phenotype associations would still exhibit some degree of spatial smoothness. We anticipate this may be a problem for methods whose null distributions involve various forms of spatial randomization or permutation. Examples of null brain-phenotype association maps from a single sub-sample from our simulations are presented in Appendix C (Figure C.1).

For NEST and FastGSEA, we compute our “observed” test statistic under *H*_0_ using the null association maps just described, and then apply the steps from Figure 2 for each network to obtain an enrichment score.

While the observed ES is identical for NEST and FastGSEA (for a given network and participant sample), their null distributions are quite different. In FastGSEA, we repeatedly permute *locations* in the network map before re-computing each ES to form the null distribution. In NEST, we permute *participants*, obtain an entirely new association map (i.e., new set of *T* (*v*)s), and then re-compute an ES to form the null distribution. For the spin test, we compute an “observed” similarity metric (Pearson correlation), which compares the same “observed” null map of brain-phenotype associations used for NEST and FastGSEA to a binary network partition map. The null distribution consists of correlations between the “observed” null map and rotations of the network map.

For each brain-phenotype association, each network, each *N*_*sub*_, and each method, we repeat these steps for 1000 simulations, estimating *p*-values based on 999 permutations (or rotations, for the spin test) per simulation. Type I error rates in each setting are estimated as the proportion of simulations yielding a *p*-value smaller than the nominal *α* = 0.05.

##### Power simulations

We estimate power by applying NEST within 1000 sub-samples of each *N*_*sub*_ (from un-permuted data). We estimate power as the rate of rejecting the null hypothesis at the *α* = 0.05 level.

(Note that we refer to this quantity as “power” for simplicity; however, in settings where the null hypothesis is true, it should resemble type I error).

As our type I error simulations will show, neither FastGSEA nor the spin test control type I error levels in the context of this realistic simulation design. Therefore, we do not consider these methods in our power simulations or in subsequent analyses.

#### 2.3.3 Data analysis

Following our simulation studies, we apply NEST to analyze the same brain-phenotype associations using data from the full PNC sample. Again, we quantify associations at the vertex level for cortical thickness and *n*-back with age, sex, and the interaction between age and sex using the same approach described in Section 2.3.1 above. In this analysis, *p*-values are estimated based on *K* = 4, 999 permutations.

We report un-adjusted *p*-values for all seven networks and indicate which remain statistically significant after controlling the false discovery rate (FDR) at *q <* 0.05. In our adjustment for multiple comparisons, we conservatively account for testing across both modalities, three brain-phenotype associations, and seven networks, for a total of 42 tests.

## 3 Results

### 3.1 Simulation results

As described in Section 2.3.2, our type I error simulation studies evaluate the rate at which each method (NEST, spin, and FastGSEA) reject their respective null hypotheses in settings where our proposed null hypothesis, as defined in Section 2.1, is true. Results are presented in Figure 3, with different color points used for each method (blue for NEST, pink for spin, and yellow for FastGSEA) and different shades used to distinguish between sample sizes (darker shades indicate larger sample size).

**Figure 3.**
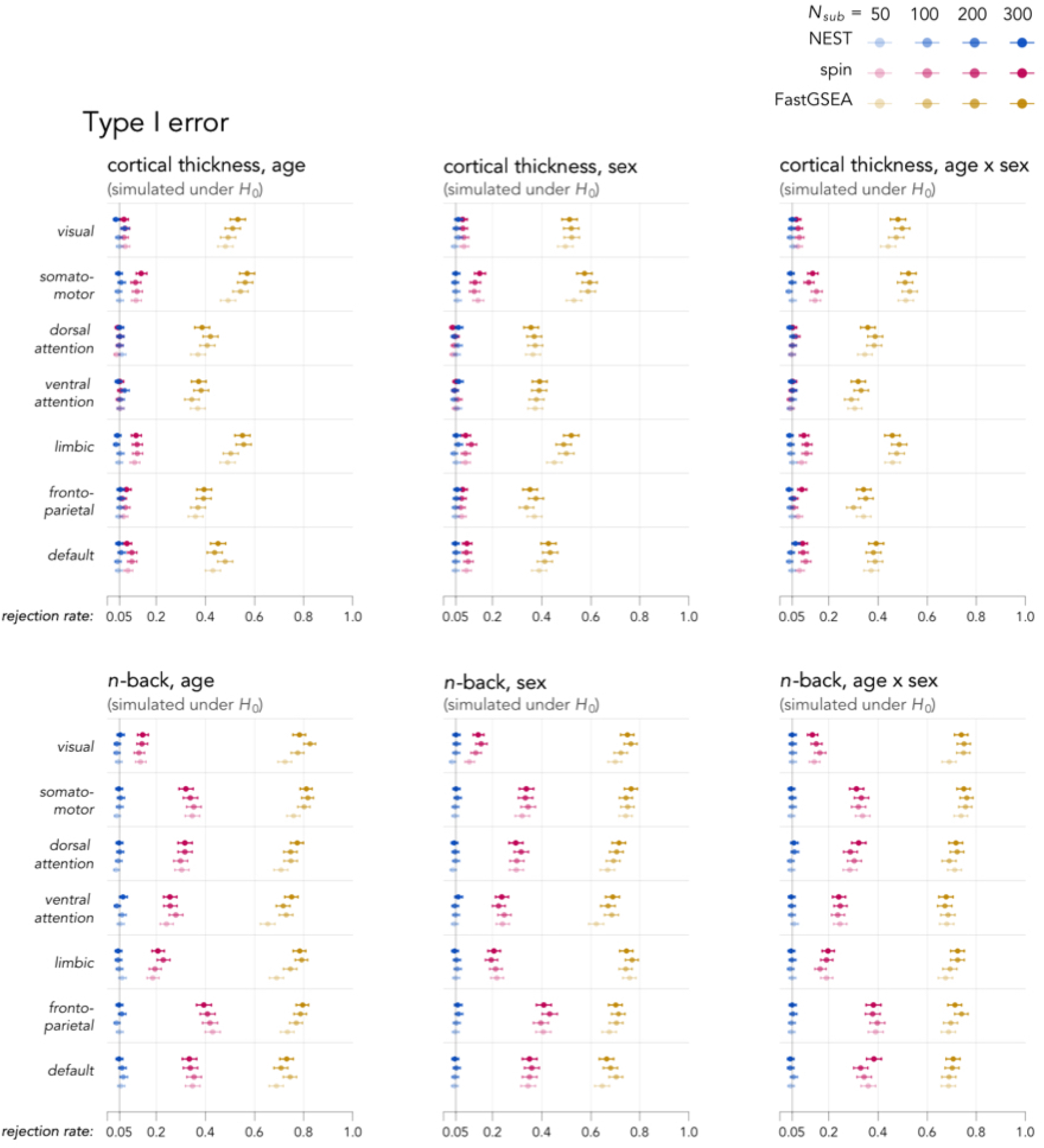
Comparison of type I error levels of our proposed method (NEST), the spin test (Alexander-Bloch et al. 2018), and FastGSEA (Korotkevich et al. 2016; Park et al. 2018). 95% binomial confidence intervals are shown with segments surrounding each point (may not be visible for especially small intervals). We find that NEST controls type I error levels at the nominal level (*α* = 0.05) in simulation studies involving all brain-phenotype associations, all networks, and all sample sizes. In contrast, neither the spin test nor FastGSEA control type I errors, suggesting these methods may not be reliable in tests of network enrichment. We speculate that this may be due to inherent spatial smoothness of null brain-phenotype association maps, even in the absence of network enrichment. Examples of simulated null association maps involving the *n*-back task are shown in Figure C.1 of Appendix C.

NEST successfully controls type I error levels at the nominal (*α* = 0.05) across all simulated null brain-phenotype associations, all sample sizes, and all seven networks. In contrast to NEST, neither the spin test nor FastGSEA controls type I error levels. For the spin test, type I error rates range from 10.5% [95% binomial CI: 0.9-12.6%] to as high as 43.2% [40.2-46.3%] for null simulations involving the *n*-back task and from 3.7% [2.7-5.1%] to 14.9% [12.8-17.3%] for those involving cortical thickness. Although the spin test does control type I error levels in certain situations (e.g., simulated nulls involving cortical thickness-age, sex, and age×sex associations in the ventral attention network), more often, we reject the null at a rate that exceeds the nominal level. FastGSEA also fails to control false positive rates, with estimates ranging from 29.1% [26.4-32.0%], at best, to 82.6% [80.1-84.8%], at worst. Given these findings, we exclude both FastGSEA and the spin test from subsequent results, as these methods’ proneness to inflated false positive findings calls into question their interpretation.

In Figure 4, we present the statistical power of NEST after repeatedly applying the method in each randomly selected sub-sample from the full observed (i.e., not permuted) PNC sample. As noted before, the null hypothesis may be either true or false, but for simplicity, we refer to to the rate of rejecting *H*_0_ as power. For cortical thickness-age effects, NEST’s power is highest in the default, frontoparietal, and dorsal and ventral attention networks. In the default network, we estimate power at 100% [99.5-100%] when *N*_*sub*_ = 200 or 300, 98.3% [97.3-99.0%] when *N*_*sub*_ = 100, and 83.4% [81.0-85.6%] when *N*_*sub*_ = 50. In the dorsal attention network, power is estimated at 100% [99.5-100%] when *N*_*sub*_ = 300, 99.5% [98.8-99.8%] when *N*_*sub*_ = 200, 91.2% [89.3-92.8%] when *N*_*sub*_ = 100, and 70.3% [67.4-73.1%] when *N*_*sub*_ = 50. In the frontoparietal and ventral attention networks, the reduction in power associated with decreasing sample sizes is more notable.

**Figure 4.**
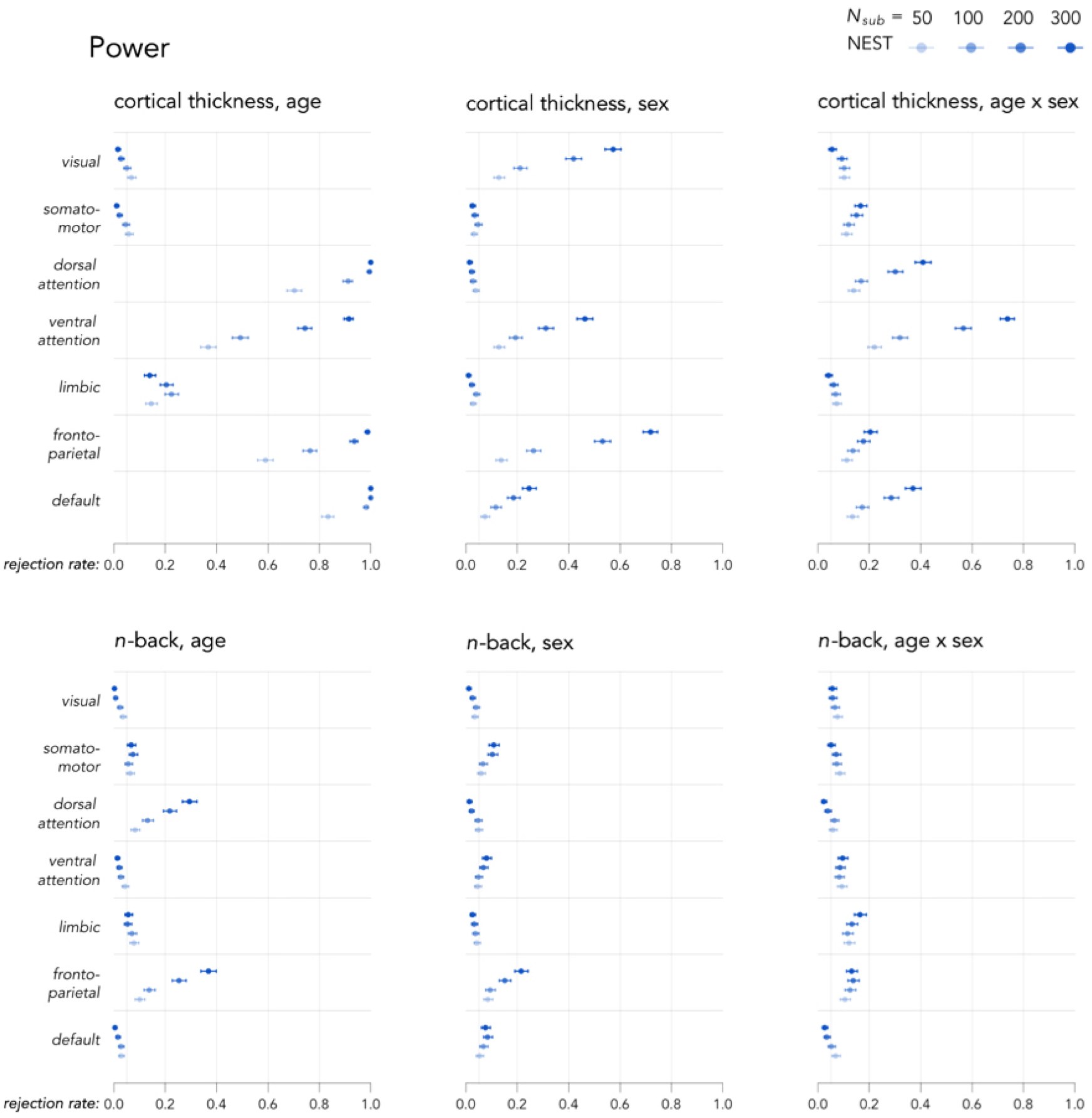
Power of NEST based on data-driven simulation studies. 95% binomial confidence intervals are indicated with line segments surrounding each point (not visible for especially small intervals).

For the frontoparietal network, power exceeds 90% when *N*_*sub*_ *≥* 200, but declines to 76.4% [73.7-78.9%] when *N*_*sub*_ = 100 and to 59.0% [55.9-62.0%] when *N*_*sub*_ = 50. For the ventral attention network, power exceeds 90% when *N*_*sub*_ = 300 but declines to 74.4% [71.6-77.0%] when *N*_*sub*_ = 200, to 49.2% [46.1-52.3%] when *N*_*sub*_ = 100, and to 36.7% [33.8-39.7%] when *N*_*sub*_ = 50.

For enrichment of associations involving the *n*-back task, it appears that larger sample sizes (beyond those considered these simulations) may be required to detect enrichment of associations where we would expect them (for instance, age effects in the frontoparietal network).

### 3.2 Data analysis results

In Figure 5, we present results from applying NEST in the full PNC cohort to evaluate the same brain-phenotype associations and networks as in our simulation studies. Unadjusted *p*-values are reported, with bolded text in Figure 5 indicating results that remain statistically significant after FDR control.

**Figure 5.**
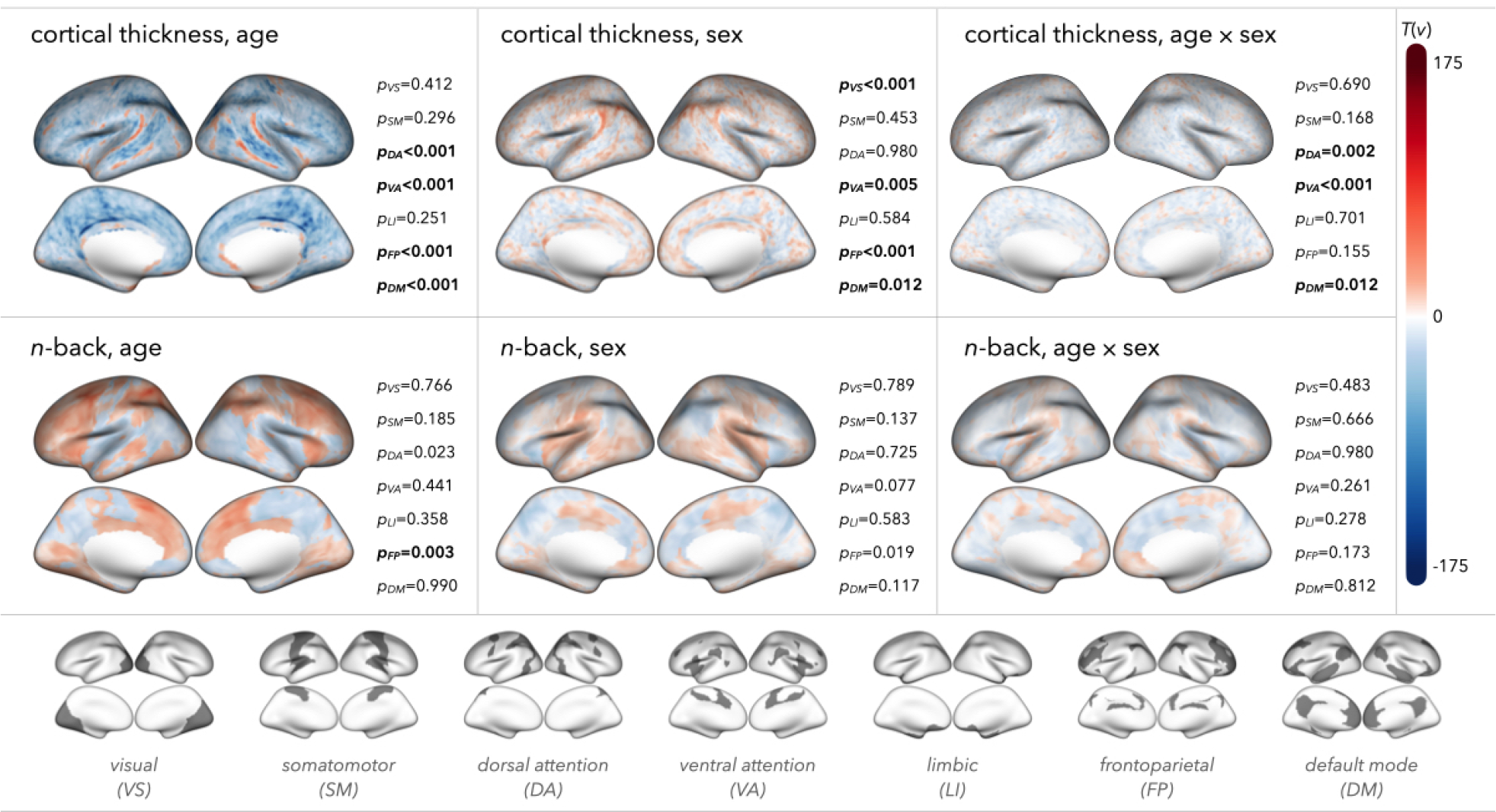
Results from application of NEST to evaluate network enrichment of associations between cortical thickness or *n*-back with age, sex, or age × sex interactions. For each setting, we map association statistics (signed multivariate Wald statistic, as described in Section 2.3.1 and Appendix B) across the cortical surface. Unadjusted *p*-values are reported for each of Yeo et al. (2011)’s seven networks, with bold text indicating results that remain statistically significant after controlling the false discovery rate (FDR) at *q <* 0.05 across both brain measurements (cortical thickness, *n*-back), all three phenotypes (age, sex, age×sex), and all seven networks.

To guide in the interpretation of Figure 5, we note that for maps involving associations with age, positive values of *T* (*v*) (shown in shades of red) indicate areas where either cortical thickness or *n*-back activation is typically found to increase (nonlinearly) with age, while negative values (shades of blue) are where we estimate the brain measurement to decrease with age. For cortical thickness-age associations, we find significant enrichment of nonlinear age effects in the default, frontoparietal, ventral attention, and dorsal attention networks. For *n*-back-age associations, we find significant enrichment in the frontoparietal network.

For interpreting spatial maps of associations with sex, positive values of *T* (*v*) indicate locations where we find either cortical thickness or *n*-back activation to be higher for females compared with males, while negative values of *T* (*v*) indicate locations where the brain measurement is found to be lower for females. After FDR correction, we find significant enrichment of sex differences in cortical thickness for the visual, ventral attention, frontoparietal, and default mode networks. We find no significant enrichment of sex effects in *n*-back activation in any networks.

For network enrichment of age × sex interactions, positive values of *T* (*v*) are interpreted as regions where nonlinear age effects tended to be higher among females compared to males, while negative values indicate locations where nonlinear age effects were generally found to be lower among females. We find significantly enriched age × sex effects on cortical thickness within the dorsal attention, ventral attention, and default mode network. We do not find significant enrichment of these effects on *n*-back in any of the seven networks.

## 4 Discussion

Interpreting brain-phenotype associations within functional networks is a common practice in neuroimaging studies, as these networks can help to link individual differences to canonical neural processes. However, existing methods for testing network specificity have fallen short by relying on subjective interpretations or strong statistical assumptions. In this article, we proposed a new framework, called NEST, for testing network enrichment.

NEST is based on Gene Set Enrichment Analysis (GSEA), a method first described by Subramanian et al. (2005), which has been widely used in genomics research for nearly two decades. Translating GSEA to the neuroimaging setting, we estimate local brain-phenotype associations to construct a spatial map of these associations. We then compute an enrichment score (ES) for a network of interest, quantifying the degree to which vertices where the brain measurement is strongly associated (either positively or negatively) with the phenotype occur within versus outside the network of interest (Figure 2). Finally, we construct a null distribution by re-computing the ES after repeatedly permuting the original data across participants and re-computing local brain-phenotype associations. We estimate a *p*-value by comparing the observed ES to this permutation-based null distribution.

Until recently, researchers often relied on ad-hoc methods to make claims about the specificity of brain-phenotype associations to functional networks. For instance, visualizing a brain map of association statistics next to a network partition map (e.g., Yeo et al. (2011), Schaefer et al. (2018)), researchers would comment on the appearance of overlap in brain regions with especially large statistics with networks of interest. Another approach would be to note the percentage of vertices or voxels with strong associations overlapping with networks of interest.

The spin test (Alexander-Bloch et al. 2018) transformed the way we go comparing brain maps, and there are numerous examples of researchers using it to assess network enrichment. However, while this method has become increasingly popular, it is questionable whether the null hypothesis it tests—that there is no spatial alignment between two maps—is an appropriate translation of the scientific questions it has often been used to answer (e.g., are associations in the frontoparietal network especially strong?). Results from our simulation studies indeed call into question the appropriateness of the spin test for testing these null hypotheses. In particular, we found inflated type I error rates in simulated settings with no enrichment, and yet the spin test still rejected its null hypothesis. In other words, significant spatial alignment (as suggested by the spin test) does not necessarily reveal anything about spatial enrichment, which we conceptualize to refer to a setting where strong associations are found within a network at a disproportionate rate than outside the network. As we noted before, we speculate that this is due to the spatial smoothness present even in null association maps, such as those in Appendix C. The inherent spatial structure of a null brain map would also explain the severe type I error inflation we found for FastGSEA, an alternative adaptation of GSEA (Korotkevich et al. 2016) that was previously applied to neuroimaging data (Park et al. 2018).

Several limitations of this article present opportunities for future work. Adding to recent discussions concerning power in neuroimaging studies (Marek et al. 2022), it would be worthwhile to explore possible ways to improve NEST’s power when large sample sizes are not available or feasible. Indeed, our simulation studies revealed a noticeable reduction in power in smaller samples (Figure 4). While we quantified local brain-phenotype associations at the vertex-level and did not leverage spatial information in these association statistics, in future work, we might explore whether we can enhance signal (and therefore improve our ability to detect enrichment) in smaller-sample settings by leveraging spatial information in quantifying brain-phenotype associations (e.g., Park and Fiecas (2022)).

In addition, unlike earlier methods, NEST requires participant-level data given that participants (rather than spatial locations) are the unit of randomization. Future work should therefore consider alternative approaches that do not require participant-level data. Furthermore, while our simulations revealed inflated type I error rates for the spin test, in future work, it would be beneficial to explore a wider variety of scenarios in which the method may be valid.

While assessing network specificity of brain-phenotype associations is quite common in neuroimaging studies, existing approaches are severely limited by either subjectivity or reliance on strong assumptions. NEST is the first method, to our knowledge, that circumvents these limitations.

## Acknowledgments

This work is supported by R01MH123550 (RTS), R01NS112274 (RTS), R01MH112847 (TDS, RTS), R01MH113550 (TDS), R01MH120482 (TDS), R01EB022573 (TDS), R37MH125829 (TDS), R01MH119219 (RCG, REG), K23MH113118 (EBB), 1ZIAMH002949 (AR), R01MH123563 (S𝒩V), R01MH132934 (AAB), and the Brain and Behavior Research Foundation (EBB).

## Data and Code Availability

Neuroimaging data were acquired as part of the Philadelphia Neurodevelopmental Cohort (PNC). PNC data are publicly available in raw format at https://www.ncbi.nlm.nih.gov/projects/gap/cgi-bin/study.cgi?study_id=phs000607.v3.p2. An R package for implementing NEST is under development and will be made available before publication.

## Author Contributions

**Sarah M. Weinstein**: conceptualization, methodology, software, writing (original draft), reviewing, and editing. **Simon N. Vandekar**: supervision, methodology, writing, reviewing, and editing. **Aaron F. Alexander-Bloch**: methodology, software, writing, reviewing, and editing. **Armin Raznahan**: methodology, writing, reviewing, and editing. **Mingyao Li**: methodology, writing, reviewing, and editing. **Raquel E. Gur**: data collection, data processing, writing, reviewing, and editing. **Ruben C. Gur**: data collection, data processing, writing, reviewing, and editing. **David R. Roalf** : data collection, writing, reviewing, and editing. **Min Tae M. Park**: conceptualization, writing, reviewing, and editing. **Mallar Chakravarty**: conceptualization, writing, reviewing, and editing. **Erica B. Baller**: data processing, writing, reviewing, and editing. **Kristin A. Linn**: supervision, methodology, writing, reviewing, and editing. **Theodore D. Satterthwaite**: supervision, conceptualization, data collection, data processing, writing, reviewing, and editing. **Russell T. Shinohara**: supervision, conceptualization, methodology, writing, reviewing, and editing.

## Appendix A Data collection and pre-processing

In this study, we use data from the Philadelphia Neurodevelopmental Cohort (PNC), a large-scale research study of children, adolescents, and young adults led by researchers at the University of Pennsylvania and Children’s Hospital of Philadelphia (Satterthwaite et al. 2014; Satterthwaite et al. 2016). Participants (*N* = 1, 601) underwent MRI scans in a Siemens TIM Trio 3 tesla machine with a 32-channel head coil. Imaging sequences and parameters are detailed in previous work (Satterthwaite et al. 2014; Satterthwaite et al. 2016). After exclusion criteria (see Weinstein et al. (2021)), we include *N* = 911 participants in our analysis of cortical thickness data and *𝒩* = 1, 018 participants in our analysis of *n*-back activation.

Structural MRI scanning protocols involved a magnetization-prepared, rapid-acquisition gradient echo (MPRAGE) T1-weighted image with 0.9 × 0.9 × 1.0 mm voxel resolution, and image quality was rated by three experienced analysts (Rosen et al. 2018). Using FreeSurfer (version 5.3), we converted T1-weighted images to cortical surface data (including template registration, intensity normalization, and inflation of cortical surfaces to the fsaverage5 template). We quantified cortical thickness as the minimum distance (in mm) between the pial and white matter surfaces (Dale et al. 1999).

For functional imaging during the *n*-back task, a single-shot-interleaved multi-slice, gradient-echo, echo planar imaging sequence with voxel resolution of 3 × 3 × 3 mm was used. We used the eXtensible Connectivity Pipeline (XCP) Engine for to preprocess the data and mitigate noise induced by head motion (Ciric et al. 2018). During the *n*-back task, participants viewed geometric stimuli and were instructed to press a button if a present stimulus matched the *n*th last one they viewed. In this study, we examined *n*-back activation maps quantifying local percent changes in brain activation between the 2-back and 0-back (i.e., pressing button regardless of relationship between current and previous stimuli) task (Ragland et al. 2002).

## Appendix B Metric for quantifying brain-phenotype associations

In this study, we quantify location (*v*)-specific brain-phenotype associations, *T* (*v*), using a signed multivariate Wald statistics. The following steps describe how we obtain this metric.

1. We consider the model of brain measurement **x**(*v*) (as a function of phenotypes or covariates) at each image location (*v*), as presented in presented in Equation (9):

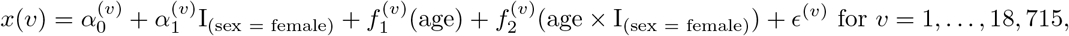

Where 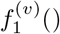 allows for nonlinear age effects 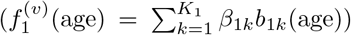, and 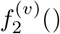 allows for nonlinear age effects that may differ by sex 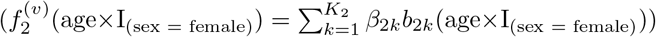 at location *v*.
2. Let 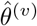 denote a vector of all the coefficients estimated in (1) for a given *v*, including 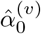 (mean), 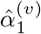 linear age effect), and all the 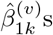 (nonlinear age effects) and 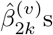 (nonlinear age × sex effects). Let 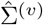 denote the variance/covariance matrix of 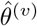 . Let *Y* denote the design matrix for the GAM in (1)and 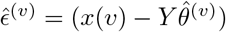(residuals).

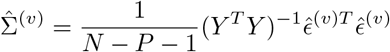

where *P* = 2 + *K*_1_ + *K*_2_ (number of parameters estimated in (1)). Note: the dimension of 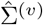 is *P* × *P*, with each row/column corresponding to a different parameter. In the next step, we use subscripts to denote sub-sets of the vector 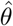 and matrix 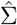 corresponding to specific parameters being captured in the Wald statistic.
3. The multivariate Wald statistics for testing the absence of sex, age, and age × sex effects are defined as follows:

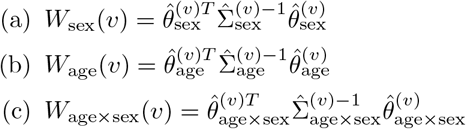
4. While the GAM considered above captures nonlinear associations, we still would like to approximate the overall trend in direction of each association with a (positive/negative) sign. For this, we use coefficient estimates from vertex-level multiple regression models. For age and sex effects, we fit the following model with no interaction term:

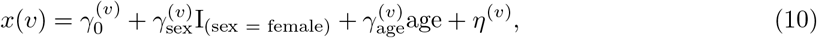

so that the signs of the coefficient estimates, sign 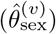 and sign 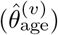 reflect both the marginal and interaction effects. To obtain the sign for the age × sex interaction, we again fit the model in Equation (10) but with an interaction term included 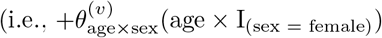 in Equation (10)).
5. Our association metric for a given brain-phenotype association at location *v, T* (*v*), is the product of the multivariate Wald statistic obtained in step (3) and the sign of the corresponding estimated coefficient from the linear model in step (4).

## Appendix C Supplementary results

**Figure C1.**
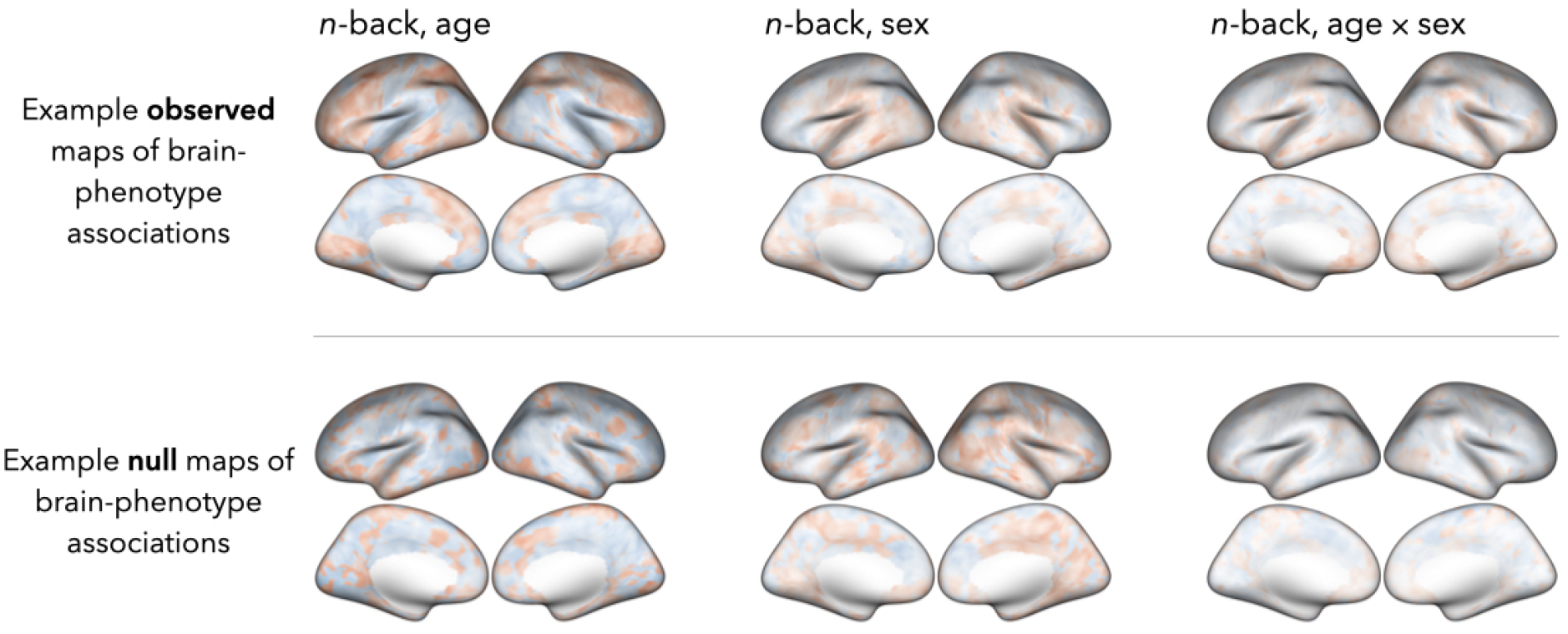
Example maps of *n*-back-age, sex, and age × sex associations from one simulation setting (with *N*_*sub*_ = 300). Observed vertex-level statistics are estimated after randomly selecting 300 PNC participants from the full sample. Null statistics are estimated after first permuting the full P𝒩C sample and then randomly selecting 300 participants from the permuted data. The null maps appear to exhibit spatial smoothness and structure, which may explain the type I error rate inflation we observe for both the spin test and FastGSEA.

